# Trends in hoaxes of academic communication

**DOI:** 10.1101/2024.11.17.624043

**Authors:** Zen Faulkes

## Abstract

Academic journals use peer review to weed out false information, but peer review and other editorial processes are normally confidential. Therefore, individuals sometimes create hoaxes to test whether editorial processes are as robust as they are claimed, or whether they are done at all. This article tracks the occurrence of hoaxes aimed at scholarly publishers and academic conferences since 2000. Since 2009, successful hoaxes usually appeared at a year of one or more a year, usually motivated by academics or journalists exposing so-called “predatory” journals. The apparent rise in the number of hoaxes reflects a lack of transparency in editorial processes at both legitimate and “predatory” journals. Reaction of academic communities to hoaxes varies widely depending on the perceived intent of the target of the hoaxes and whether the hoax demonstrates what the hoaxer claims.

## Introduction

Hoaxes have been part of the scholarly landscape for centuries (MacDougall 1940; Walsh 2006; Young 2017). Experts and academics are particularly tempting targets for hoaxes, because they are often perceived as prone to overestimating their own intelligence. Many people can appreciate the irony of Sherlock Holmes’s creator, Sir Arthur Conan Doyle, being fooled by faked pictures of fairies (Conan Doyle 1922; Young 2017). Moreover, an expert who is fooled might declare a hoax to be genuine, which can help a hoax reach a wider audience (Young 2017). Some examples of hoaxes targeting researchers include the Piltdown man hoax of faked human fossils (Weiner et al. 1953; Straus 1954; Russell 2003; Russell 2012) and the fake psychics of the Project Alpha hoax (Randi 1983b; Randi 1983a).

Hoaxes perpetrated by academics were rarer. Alan Sokal’s hoax (Sokal 1996) received much attention and analysis (Editors of Lingua Franca 2000) for at least two reasons. First, it was an academic criticizing the practices of other academics who were acting in good faith. Second, it was perceived as an extraordinary “one off” event. Sokal’s hoax created a model for others to follow, and there has been a string of academic hoaxes since. Previous examinations of hoaxes of academics (Djuric 2015; Al-Khatib and Teixeira da Silva 2016) have analyzed small numbers of hoaxes as case studies. My goal is to describe larger patterns of hoaxes over many years. As such, this paper includes some hoaxes that have been previously documented and discussed in the literature (e.g., (Djuric 2015)). Other hoaxes are known from mainstream media or social media but have not previously been documented in academic literature.

Hoaxes should be studied for several reasons. First, hoaxes have the potential to do harm. Most discussion of hoaxes has focused on the potential to erode trust. Another potential for harm is that a hoax that is seen by its creator as targeting a predatory journal may in fact be an legitimate journal that is under resourced or inexperienced. Second, those on the receiving end of hoaxes have vested interests in concealing that they have been hoaxed. Those who create hoaxes may also have vested interests in downplaying their hoaxes if they are controversial. Thus, studying hoaxes creates a more permanent record of these events in the literature, which are important in holding people accountable.

## Methods

For the purposes of this paper, a hoax is creating and disseminating information that the creator knows is untrue but that, in the opinion of the creator, believes could be recognized as deception with due diligence. The latter qualifier is important. A deception that the perpetrator does not expect people to see as untrue is better described as fabrication or falsification. Fabrication and falsification are meant to go undetected, ideally indefinitely. Thus, for the purposes of this paper, the intention of the creator matters (*contra* Young (2017), who does not consider intention to be important in determining what is a hoax). Likewise, the profession of the creator does not matter (*contra* Al-Khatib and Teixeira da Silva (2016), who distinguished “stings” from “hoaxes” by profession: stings were create by journalists, hoaxes by anyone else).

Similarly, some of the hoaxes in this paper were categorized as “irony” by Al-Khatib and Teixeira da Silva (2016), although they did not define the term. Some of their examples might be described as humour, spoofs, or parodies. (Ronagh and Souder 2015). Articles intended as humour or parodies by their authors can be misread by people as being real (Asimov 1972; Ronagh and Souder 2015). As noted above, the intention of the creator matters. The creator of a hoax expects that someone might believe it is real if they are not paying attention. The creator of a spoof does not expect anyone to believe it is real. For example, someone who tells a joke that begins, “A duck walks into a bar and asks for a drink,” does not expect to have to explain that ducks do not, in fact, talk.

Hoaxes were collected *ad hoc* from July 2017 to June 2020. This may not be a comprehensive account of academic publishing hoaxes in that time but should be a large enough sample to be representative of patterns of hoaxes.

I included hoaxes that were: created after the year 2000; targeted at an academic venue (e.g., journal or conference); accepted for publication by the venue; then publicly revealed as untrue (i.e., material in which the author or authors claimed the work was genuine was excluded, even if later shown by others to be false). Multiple targets that were revealed at once were considered as a single hoax (Al-Khatib and Teixeira da Silva (2016) treated hoaxes similarly).

For example, Bohannon (2013) and Norddeutscher Rundfunk et al. (2018) submitted hundreds of manuscripts to journals as part of a single, cohesive investigation, and both were considered single hoaxes. As described above, I omitted articles where there was no intention to deceive either the editorial staff or the readers. For example, (The Study of Maternal Child Kissing (SMACK) Working Group 2015) was not included because authors and editors believed the readers would be in on the joke (Al-Khatib and Teixeira da Silva 2016; Oransky and Marcus 2016).

I collected both the full text of the hoaxed article when available and the announcement(s) revealing the hoax. I archived the text of hoaxes in Faulkes (2020b) and the resulting data in (Faulkes 2020a). I analyzed the text of the hoax document for indications that the content was untrue (e.g., unintelligibility, humour, profanity).

To test the hypothesis that papers passed peer review because of the writing was complex (pseudo-sophistication), I converted the English texts from PDFs to plain text and calculated six standard readability scores (Flesch Kincaid Reading Ease, Flesch Kincaid Grade Level, Gunning Fog Score, SMOG Index, Coleman Liau Index, and Automated Readability Index) for the main body of the text (i.e., excluding title, author information, and references) using readbilityformulas.com website (https://readabilityformulas.com/freetests/six-readability-formulas.php). Three hoaxes were omitted from readability analyses: one that was written in German (Schulte et al. 2016) and two were not organized in typical paragraph structure (Keulemans 2017; Sorokowski et al. 2017).

The announcements of hoaxes were analyzed for what prompted the hoax, the identity of the hoaxer, the target(s) of the hoax, the number of papers in the hoax, the outcomes of the hoax (e.g., accepted versus rejected hoax papers). Given the variation in how the hoaxes were disclosed (ranging from extensive essays by the hoaxers themselves to short blog posts by people not involved in the hoax), information about the details of the hoaxes were not necessarily consistently disclosed. For example, some hoaxers described the details of article processing fees but others did not.

To test whether hoaxed articles would remain available and potentially cause confusion to naïve readers, I checked all articles for assigned digital object identifiers (DOIs). Using a standard web browser, I checked whether I could located the original hoax article using the DOI-based link (“https://doi.org/“ followed by the DOI; e.g., https://doi.org/10.6084/m9.figshare.5248264, which leads to (Faulkes 2020b)). I conducted this check for working DOI links on 9 June 2020.

While a comprehensive survey of reactions to the hoaxes presented was outside the scope of this project, responses to hoaxes were collected on an *ad hoc* basis and archived in Faulkes (2020b).

## Results

Twenty-seven hoaxes were gathered (Table 1).

**Table 1.**
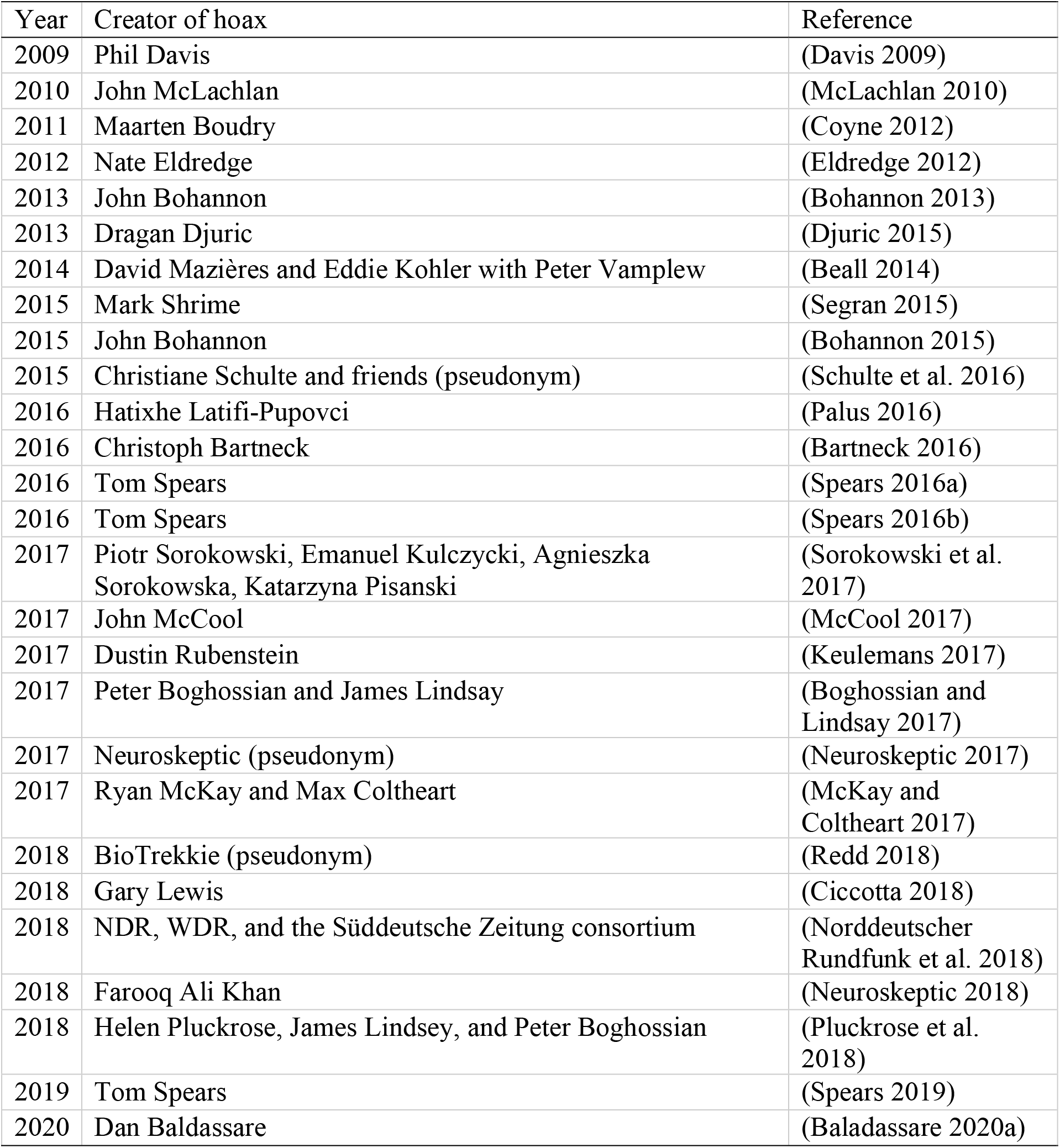
List of hoaxes included in the paper.

The frequency of hoaxes increased over time (Figure 1). There was one hoax a year from 2009 to 2012 and in 2019, but multiple hoaxes (two to six) every year from 2013 to 2018.

**Figure 1.**
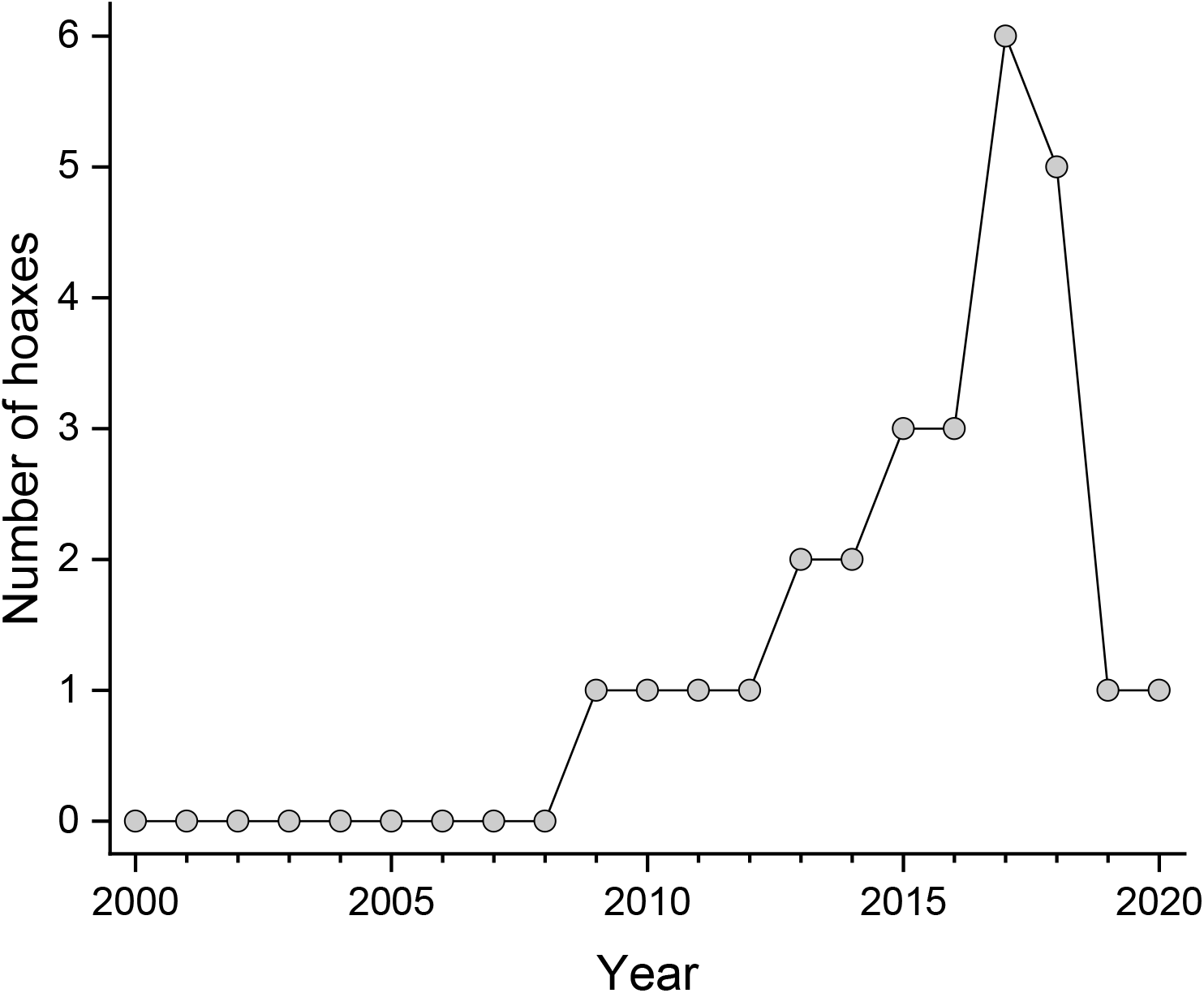
Number of academic hoaxes per year from 2000 to 2020.

**Figure 2.**
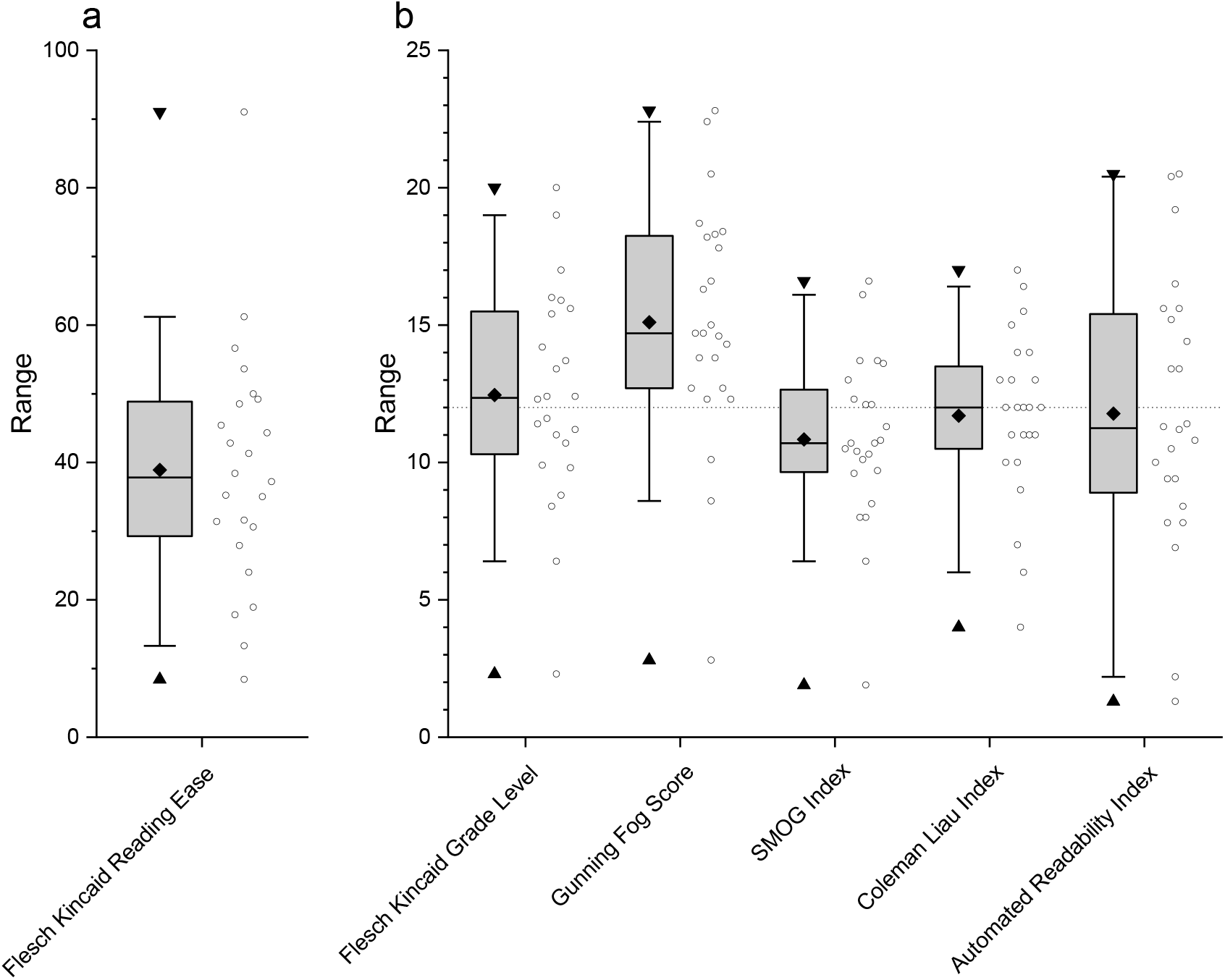
Readability scores for text of hoaxed articles. (a) Flesch Kincaid Reading Ease, where higher scores indicate greater readability. (b) Scores for Flesch Kincaid Grade Level, Gunning Fog Score, SMOG Index, Coleman Liau Index, and Automated Readability Index. All indices roughly correspond to reading grade levels. A score of 12 corresponds to high school graduate reading level, shown by dotted line. A score above 12 corresponds to university level reading. Diamond = mean; horizontal line = median; box = 50% of data; whiskers = 95% of data; triangles = minimum and maximum.

The most common form of hoax was a faked journal article (23/27, 85.2%). The remaining hoaxes were faked conference abstracts (4/27, 14.8%) (McLachlan 2010; Coyne 2012; Bartneck 2016; Schulte et al. 2016), a faked biography of a scientist (1/27, 3.7%) (Sorokowski et al. 2017), and a faked journal (1/27, 3.7%) (Keulemans 2017). Numbers do not add up to 100% because one hoax consisted of both a conference presentation and a publication (Schulte et al. 2016).

Most hoaxes (21/ 27, 77.8%) were perpetuated by academics. The rest (6/27, 22.2%) were created by journalists investigating academic publishing or academic journalism (Bohannon 2013; Bohannon 2015; Spears 2016a; Spears 2016b; Spears 2019). Seven hoaxes (7/27, 25.9%) were created by people involved in multiple hoaxes (Bohannon 2013; Bohannon 2015; Spears 2016a; Spears 2016b; Boghossian and Lindsay 2017; Pluckrose et al. 2018; Spears 2019).

The scope of hoaxes varied widely. Most hoaxes (17/27, 63.0%) were created by academics who wrote a single paper or abstract that was submitted to a single journal or conference. At the other end of the spectrum, two hoaxes submitted manuscripts to more than 300 journals (Bohannon 2013; Sorokowski et al. 2017). One hoax submitted fewer papers, but was part of a large investigation run by consortium of journalists investigating predatory journals generally (Norddeutscher Rundfunk et al. 2018).

The most common motivation for the hoax (21/27, 77.7%) was to expose predatory journals or predatory publishers. Multiple hoaxers (8/27, 29.6%) specifically mentioned unsolicited emails (also known as “spam”) as a prompt for their hoax (McLachlan 2010; Stromberg 2014; Bartneck 2016; Spears 2016b; Keulemans 2017; McKay and Coltheart 2017; Lewis 2018a; Baladassare 2020a). The intent of these hoaxes was to show that some venues publish anything for money, but several hoaxers (4/21, 19.0%) negotiated with publishers to reduce the price (Keulemans 2017; Ciccotta 2018; Redd 2018; Baladassare 2020b). In two cases, the journal waived the article processing fee entirely (Ciccotta 2018; Lewis 2018b; Baladassare 2020b). Further, McKay and Coltheart (2017) noted that some publishers ran their submission through plagiarism detectors.

In some cases (5/27, 18.5%), the hoaxers viewed their target as fields of scholarship rather than individual journals or publishers. These generally targeted disciplines in the humanities (Coyne 2012; Schulte et al. 2016; Boghossian and Lindsay 2017; Pluckrose et al. 2018). Only one hoax (1/27, 3.7%) might be seen as targeting a scientific or technical discipline writ large. McLachlan (2010) wrote his conference abstract “parodied the absurdity and credulity of so-called integrative medicine.”

One hoax was conducted as part of a larger effort to show shortcomings in science journalism (Bohannon 2015). Although the publication of the paper exposed a predatory publisher, that was a means, not an end.

Most hoaxes were written by the creators (19/27, 70.4%). Four hoaxers (4/27, 14.8%) generated text randomly, resulting in text that was grammatically correct but nonsensical. Three hoaxes (3/27, 11.1%) used plagiarized text.

The average readability scores for hoax texts were low, with most scores indicating most papers required about a Grade 11 or Grade 12 (i.e., high school graduate) reading level. The variation in readability was substantial, however. A paper titled “Get me off your fucking mailing list” was rated the most readable by all six scores used (Beall 2014). The paper consists of the title repeated throughout the text, with almost no other words. Likewise, a paper titled “The conceptual penis as a social construct” (Boghossian and Lindsay 2017) was rated the least readable by all six scores.

The clues intended to show that a contribution should be rejected were often humorous. A popular format was to write the entire paper as an extended reference to, or parody of, a pop culture movie or television episode: e.g., *Star Wars* (Neuroskeptic 2017), *Seinfeld* (McCool 2017), *Star Trek: Voyager* (Redd 2018), and *Rick and Morty* (Neuroskeptic 2018). Some were more subtle, inserting short references to pop culture, such as listing the Warner Brothers cartoon character Yosemite Sam as an author (Spears 2019), listing the *Sesame Street* character Big Bird in the acknowledgements (Baladassare 2020a), or referencing the German TV show *Kommissar Rex* (Schulte et al. 2016).

Proving that even academics are not immune to the joys of bathroom humour, hoaxed papers included profanity (Safi 2014) and / or references to male genitals (Boghossian and Lindsay 2017; Pluckrose et al. 2018), buttocks (McLachlan 2010; Djuric 2015; Keulemans 2017), or feces (Davis 2009).

The hoaxes were most often revealed online via blogs or social media (13/27, 48.1%). These hoaxes were often picked up by more traditional news outlets later. Traditional news outlets such as newspapers, magazines, and news websites were the other main platforms for hoaxers to reveal their work (12/27, 44.4%). The remaining three hoaxes (3/27, 11.1%) were revealed in academic journals, although these journals also featured news and views in addition to academic articles.

About 40% of hoaxes (11/27, 407%) had at least one article that was assigned at least one DOI. Some hoaxes had multiple papers that were assigned DOIs, for 15 DOIs total. These were evenly split between leading to retraction notices (7/15, 46.7%) and receiving error messages that the article could not be found (7/15, 46.7%). One DOI (1/15, 6.7%) was active but lead to a completely different article than the original hoax article.

## Discussion

Academic hoaxes in the twenty-first century fall into two broad categories. First, there are hoaxes meant to expose bad publishing practices in science, technology, engineering, and math (STEM disciplines). Second, there are hoaxes meant to impugn humanities for poor scholarship.

The STEM hoaxes are an unintended consequence of the push for open access in academic publishing in the 2000s and 2010s. The success of open access journals like *PLOS ONE* normalized the use of article processing charges: payment by authors of accepted articles. While charges to authors existed long before the 2000s (usually referred to as “page charges”), the use of article processing charges in turn inspired the creation and proliferation of so-called “predatory” journals: journals that charge authors, place articles on websites, but did none of the editorial quality control that characterized legitimate academic journals. (The phrase “predatory journal” is not ideal, because there is little agreement on how to define a “predatory” journal or publisher (Grudniewicz et al. 2019), but it is the best known phrase for exploitative journals.) The first hoax in this study (Davis 2009) predates the term “predatory journal” or “predatory publisher” (coined by librarian Jeffrey Beall in 2010; (Butler 2013)), but it is probably not coincidental that hoaxes increased after the term “predatory publisher” appeared in *Nature*, a widely read weekly science journal (Beall 2012) as academics became more widely aware of the existence of predatory journals and the problems posed by them. The decline of hoaxes after 2017 may be because the problem of predatory journals was better known by then, and fewer people felt the need to demonstrate that they exist.

Hoaxes of predatory journals have revealed features that might not otherwise be obvious: that many of the people behind predatory journals are apparently incompetent at extracting money. It is not clear why multiple hoaxers were able to negotiate reduced article processing fees, two of whom had article processing fees waived (Lewis 2018b; Baladassare 2020a).

Likewise, it is unclear why predatory publishers would run plagiarism checks (McKay and Coltheart 2017). One possibility is that even a predatory journal needs to publish new papers continually to be seen as viable to naïve potential authors. The publishers may be willing to publish a paper for no money essentially as a form of advertising for the journal.

Investigations of predatory publishers have tended to focus on the level of quality control and the individuals who publish in them. Neither scientists nor journalists have ever investigated the individuals behind predatory publishers to discover their motivations and practices.

The humanities hoaxes were meant to question the credibility of entire fields. The hoaxers targeted mainstream academic publications and conferences in the field, rather than predatory journals. In one case when a humanities hoax was published in a predatory journal (Boghossian and Lindsay 2017), it was followed by another hoax aimed at established publications (Pluckrose et al. 2018). These hoaxes usually reference the Sokal hoax (Sokal and Bricmont 1998; Editors of Lingua Franca 2000) as inspiration. Some hoaxers of predatory journals also mention Sokal (Djuric 2015), but their goals are more limited than Sokal or the other hoaxers of humanities. None of the STEM hoaxers claim that their paper was an indictment of an entire field.

The goals of hoaxers in these two categories may reflect two differences in scholarly publishing in STEM and the humanities. Hoaxes are probably not necessary to show that flawed work passes peer review in STEM subjects, because retracted articles make that point (Steen et al. 2013). There are more retractions in STEM journals than humanities journals (Figure 3 in (Ribeiro and Vasconcelos 2018); not adjusted for number or publications, however), suggesting that some key editorial practices are different in STEM and humanities. For example, retraction for errors and misconduct are more common in STEM (Decullier et al. 2013; Halevi 2020), while duplicate publication and plagiarism is a major reason for retraction in humanities (Halevi 2020). Further, retraction practices in STEM disciplines are increasing in part because of changing expectations and higher stringency (Fanelli 2013; Steen et al. 2013).

A second difference between scholarly publishing in STEM and humanities is open access publishing. The push for open access publication began in the STEM disciplines, and has been slower to take hold in humanities (Suber 2004). As open access becomes more common in humanities disciplines, it seems likely that more predatory journals will emerge to exploit scholars in those fields, and this will probably be followed by hoaxes to demonstrate that some specific humanities journals are low quality or exploitative.

Both STEM and humanities hoaxes are potentially problematic. Hoaxes are, by definition, untrue and conducted by someone with a point to make (or, less charitably, an axe to grind). Some hoaxes are controversial because they seem intended to mock or impugn scholars rather than uncover deceitful practices. Hoaxes always have the potential to erode trust necessary for scholarly publishing to work at all (Al-Khatib and Teixeira da Silva 2016; Oransky and Marcus 2016; Siebert and Schreven 2019). Nevertheless, hoaxes may have value, because they can sometimes reveal what criticism cannot. No degree of criticism of predatory publishing would have shown that people running an allegedly predatory journal would waive article processing fees or run plagiarism checks.

Further, a hoax is one way to show that editors and reviewers are doing what they claim to. There are often good reasons for confidentiality about peer review, but this means that a journal’s editorial practices are often opaque to both authors and readers. Authors and readers may be less concerned by confidentiality if they know and trust the editors and editorial board of journals. That social trust has become hard to maintain as the number of scientists, number of journals, and the size of editorial boards has expanded. It is now difficult for a working academic to know many the editors of journals in a field personally and to gain that social trust. A hoax is an experimental test of a hypothesis about publishing practices. In this sense, hoaxes may serve a valuable purpose for demonstrating that journals are taking quality control steps that they claim to be doing all the time.

Indeed, journals that are serious about quality control and transparency could make unannounced submission of fake manuscripts part of their own quality control. Unannounced inspections are an accepted means of quality control and are recognized as important for public health (e.g., restaurant and food inspections). The unannounced submission of deliberately bad articles could serve the same purpose for journals. Hoaxers could be the scholarly publishing equivalent of “white hat hackers” in computing: people who test security with the goal of finding failure points. Journals could use false manuscripts to check their editorial processes and try to address them internally.

Hoaxes seem likely to remain part of the academic publishing landscape for the foreseeable future. The study of hoaxes may become more difficult, because each new hoax renders the next hoax less notable and newsworthy, and more and more platforms exist in which to reveal hoaxes. Further, a substantial number of hoaxes vanish from publisher websites, either as an unannounced form of retraction or because the publisher goes out of business and published articles are removed from web servers. Hoaxes should continue to be recorded and studied for their insights into scholarly publishing – and maybe a few good laughs.

## References

Al-Khatib A, and Teixeira da Silva JA. 2016. Stings, hoaxes and irony breach the trust inherent in scientific publishing. Publishing Research Quarterly, 32(3): 208–219. 10.1007/s12109-016-9473-4.

Asimov I. 1972. The Early Asimov or, Eleven Years of Trying. Doubleday, Garden CIty, New York.

Baladassare D 2020a. AMAZING news folks. I submitted my paper to the prestigious Scientific Journal of Research & Reviews https://irispublishers.com/sjrr/ xand it was accepted! They formatted it for publication, including @stacyfarina’s lovely figure! Now to come up with $1,600… In: @evornithology (ed.) AMAZING news folks. I submitted my paper to the prestigious Scientific Journal of Research & Reviews https://irispublishers.com/sjrr/ and it was accepted! They formatted it for publication, including @stacyfarina’s lovely figure! Now to come up with $1,600… 3:06 PM ed.: Twitter.

Baladassare D 2020b. FOLKS. We did it. We wore them down. WE WON. This random, sketchy, blatant scam of a predatory journal decided to publish my magnum opus FOR FREE. You can now find the final, published version in all it’s glory here: https://t.co/1AEYS6G2Lv. In: @evornithology (ed.) FOLKS. We did it. We wore them down. WE WON. This random, sketchy, blatant scam of a predatory journal decided to publish my magnum opus FOR FREE. You can now find the final, published version in all it’s glory here: https://t.co/1AEYS6G2Lv. 12:01 PM ed.: Twitter.

Bartneck C 2016. iOS just got a paper on nuclear physics accepted at a scientific conference. Christoph Bartneck, Ph.D.

Beall J. 2012. Predatory publishers are corrupting open access. Nature, 489: 179. 10.1038/489179a.

Beall J 2014. Bogus journal accepts profanity-laced anti-spam paper. Scholarly OA.

Boghossian P, and Lindsay J 2017. The conceptual penis as a social construct: A Sokal-style hoax on gender studies. Skeptic.

Bohannon J. 2013. Who’s afraid of peer review? Science, 342(6154): 60–65. 10.1126/science.342.6154.60.

Bohannon J 2015. I fooled millions into thinking chocolate helps weight loss. Here’s how.: io9.

Butler D. 2013. Investigating journals: The dark side of publishing. Nature, 495: 433–435. 10.1038/495433a.

Ciccotta T 2018. Academic journal runs hoax article about conservatives’ bathroom habits.

Breitbart. Conan Doyle A. 1922. The Coming of the Fairies. Hodder & Stoughton Ltd., London.

Coyne JA 2012. A Sokal-style hoax by an anti-religious philosopher. Why Evolution is True.

Davis P 2009. Open access publisher accepts nonsense manuscript for dollars The Scholarly Kitchen.

Decullier E, Huot L, Samson G, and Maisonneuve H. 2013. Visibility of retractions: a cross-sectional one-year study. BMC Research Notes, 6(1): 238. 10.1186/1756-0500-6-238.

Djuric D. 2015. Penetrating the omerta of predatory publishing: The Romanian connection. Science and Engineering Ethics, 21(1): 183–202. 10.1007/s11948-014-9521-4.

Editors of Lingua Franca 2000. The Sokal Hoax: The Sham the Shook the Academy. Lincoln, Nebraska: University of Nebraska Press.

Eldredge N 2012. Mathgen paper accepted! That’s Mathematics!

Fanelli D. 2013. Why growing retractions are (mostly) a good sign. PLoS Medicine, 10(12): e1001563. 10.1371/journal.pmed.1001563.

Faulkes Z 2020a. Scholarly publishing hoax data. Figshare. 10.6084/m9.figshare.12413306

Faulkes Z 2020b. Stinging the Predators: A Collection of Papers That Should Never Have Been Published. Figshare. 10.6084/m9.figshare.5248264

Grudniewicz A, Moher D, Cobey KD, Bryson GL, Cukier S, Allen K, et al. 2019. Predatory journals: no definition, no defence. Nature, 576: 210–212. 10.1038/d41586-019-03759-y.

Halevi G. 2020. Why articles in arts and humanities are being retracted? Publishing Research Quarterly, 36(1): 55–62. 10.1007/s12109-019-09699-9.

Keulemans M. 2017. Tijdschrift ASSHOLE toont aan: een vakblad oprichten is veel te makkelijk. de Volkskrant. https://www.volkskrant.nl/wetenschap/tijdschrift-asshole-toont-aan-een-vakblad-oprichten-is-veel-te-makkelijk~bb608f83/.

Lewis G 2018a. 1/n. If you are an academic you probably get spammed by predatory journals rather a lot. And if you are anything like me it probably gets a little annoying sometimes. In: @Gary_Lewis1 (ed.) 1/n. If you are an academic you probably get spammed by predatory journals rather a lot. And if you are anything like me it probably gets a little annoying sometimes. 5:34 AM ed.: Twitter.

Lewis G 2018b. I submitted a hoax manuscript to a predatory journal. The finding? Politicians from the right wipe their ass with their left hand (and vice versa) - big breakthrough! Manuscript accepted w/o review. I then haggled the OA fee down to $0 - so here it is -> http://crimsonpublishers.com/pprs/pdf/PPRS.000516.pdf. In: @Gary_Lewis1 x(ed.) I submitted a hoax manuscript to a predatory journal. The finding? Politicians from the right wipe their ass with their left hand (and vice versa) - big breakthrough! Manuscript accepted w/o review. I then haggled the OA fee down to $0 - so here it is -> http://crimsonpublishers.com/pprs/pdf/PPRS.000516.pdf. 6:34 AM ed.: Twitter.

MacDougall CD. 1940. Hoaxes. Dover Publications, Mineola, New York.

McCool JH. 2017. Opinion: Why I published in a predatory journal. The Scientist. https://www.the-scientist.com/critic-at-large/opinion-why-i-published-in-a-predatory-journal-31697.

McKay R, and Coltheart M 2017. Breaking the ice with buxom grapefruits: Pratiques de publication and predatory publishing. BishopBlog.

McLachlan JC. 2010. Integrative medicine and the point of credulity. The BMJ, 341: c6979. 10.1136/bmj.c6979.

Neuroskeptic 2017. Predatory journals hit by ‘Star Wars’ sting. Neuroskeptic.

Neuroskeptic 2018. “Rick and Morty” sting predatory journals. Neuroskeptic. Discover.

Norddeutscher Rundfunk, Westdeutscher Rundfunk, and Süddeutsche Zeitung Magazin 2018. More than 5,000 German scientists have published papers in pseudo-scientific journals. Norddeutscher Rundfunk.

Oransky I, and Marcus A 2016. Fake study on moms’ kisses risked sowing confusion just for a laugh. Stat.

Palus S 2016. Sting operation forces predatory publisher to pull paper. Retraction Watch.

Pluckrose H, Lindsay J, and Boghossian P 2018. Academic grievance studies and the corruption of scholarship. Areo.

Randi J. 1983a. The Project Alpha experiment part 2. Beyond the laboratory. The Skeptical Inquirer, 8(1): 36–45. https://skepticalinquirer.org/1983/07/the-project-alpha-experiment-part-1-the-first-two-years/.

Randi J. 1983b. The Project Alpha experiment: Part 1. The first two years. The Skeptical Inquirer, 7(7): 24–33. https://skepticalinquirer.org/1983/07/the-project-alpha-experiment-part-1-the-first-two-years/.

Redd NT 2018. Fake science paper about ‘Star Trek’ and warp 10 was accepted by ‘Predatory Journals’. Space.com.

Ribeiro MD, and Vasconcelos SMR. 2018. Retractions covered by Retraction Watch in the 2013–2015 period: prevalence for the most productive countries. Scientometrics, 114(2): 719–734. 10.1007/s11192-017-2621-6.

Ronagh M, and Souder L. 2015. The ethics of ironic science in its search for spoof. Science and Engineering Ethics, 21(6): 1537–1549. 10.1007/s11948-014-9619-8.

Russell M. 2003. Piltdown Man: The Secret Life of Charles Dawson & the World’s Greatest Archaeological Hoax. Tempus, Stroud.

Russell M. 2012. The Piltdown Man Hoax: Case Closed. The History Press, Stroud.

Safi M 2014. Journal accepts bogus paper requesting removal from mailing list. The Guardian.

Schulte C, Friend, and Friend 2016. Kommissar Rex an der Mauer erschossen? Telepolis. Germany: Heise Medien.

Segran E 2015. Why a fake article titled “Cuckoo for Cocoa Puffs?” was accepted by 17 medical journals. Fast Company.

Siebert S, and Schreven S. 2019. Protean uses of trust: A curious case of science hoaxes. Wydawnictwo Uniwersytetu Łódzkiego, 9/2019(2): 216–230. 10.18778/2450-4491.09.15.

Sokal AD. 1996. A physicist experiments with cultural studies. Lingua franca, 6(4): 62–64.

Sokal AD, and Bricmont J. 1998. Fashionable Nonsense: Postmodern Intellectuals’ Abuse of Science. Picador, New York.

Sorokowski P, Kulczycki E, Sorokowska A, and Pisanski K. 2017. Predatory journals recruit fake editor. Nature, 543: 481–483. 10.1038/543481a.

Spears T 2016a. Owner of Canadian medical journals publishes fake research for cash. Ottawa Citizen. Ottawa: Postmedia.

Spears T 2016b. This ‘predatory’ science journal published our ludicrous editorial mocking its practices. Ottawa Citizen. Ottawa: Postmedia.

Spears T 2019. Predatory science journals pivot to video. Ottawa Citizen. Ottawa.

Steen RG, Casadevall A, and Fang FC. 2013. Why has the number of scientific retractions increased? PLoS ONE, 8(7): e68397. 10.1371/journal.pone.0068397.

Straus WL. 1954. The great Piltdown hoax. Science, 119(3087): 265–269. http://www.jstor.org/stable/1681349.

Stromberg J 2014. “Get me off your fucking mailing list” is an actual science paper accepted by a journal. 21 November 2014 ed.: Vox.

Suber P 2004. Promoting open access in the humanities. Society for Classical Studies.

The Study of Maternal Child Kissing (SMACK) Working Group. 2015. Maternal kisses are not effective in alleviating minor childhood injuries (boo-boos): a randomized, controlled and blinded study. Journal of Evaluation in Clinical Practice, 21(6): 1244–1246. https://onlinelibrary.wiley.com/doi/abs/10.1111/jep.12508.

Walsh L. 2006. Sins Against Science: The Scientific Media Hoaxes of Poe, Twain, And Others. State University of New York Press, Albany, New York.

Weiner JS, Oakley KP, and Le Gros Clark WE. 1953. The solution of the Piltdown problem. In Bulletin of the British Museum (Natural History), Geology. pp. 139–146.

Young K. 2017. Bunk: The Rise of Hoaxes, Humbug, Plagiarists, Phonies, Post-Facts, and Fake News. Graywolf Press, Minneapolis, Minnesota. 480 p.

